# Interpretable Local-to-Global Estimation of Brain Aging Speed From Morphological Changes Using Longitudinal Structural MRI Data^⋆^

**DOI:** 10.64898/2026.07.13.737874

**Authors:** Yuanwang Zhang, Hongming Li, Yong Fan

## Abstract

Accurate characterization of brain aging is essential for understanding cognitive decline and assessing the risk of neurodegenerative disease. Brain age estimated from cross-sectional MRI provides a snapshot of brain health relative to chronological age, and the derived brain age delta has emerged as a promising biomarker. However, brain age delta reflects cumulative effects at a single time point and fails to capture ongoing aging dynamics. Longitudinal approaches address this limitation by estimating age differences between scan pairs to derive aging speed. Nevertheless, existing methods primarily rely on intensity or texture differences between image pairs, overlook the spatial heterogeneity of aging processes, and provide limited interpretability. To overcome these limitations, we propose a novel framework that estimates brain aging speed from longitudinal deformation fields obtained via diffeomorphic registration. Instead of solely generating a single global estimate, our method produces patch-wise local aging predictions and adaptively integrates them into a unified global prediction, improving both predictive performance and interpretability. Evaluated on large-scale datasets, our approach achieves superior accuracy compared with existing methods and enhances the identification of abnormal aging patterns in diseased populations. Code is available at https://github.com/Kateridge/Morphology-AgingSpeed.

## 1 Introduction

Brain aging is a complex and long-term process characterized by gradual cognitive decline and biological changes such as weakened neuronal communication and regional brain atrophy. While normal aging follows relatively consistent trajectories, neurodegenerative diseases often induce abnormal regional atrophy patterns [19,23]. Understanding normal aging trajectories is therefore critical for elucidating neurodegenerative disease progression and enabling precise diagnosis and personalized interventions.

Brain age has emerged as a promising imaging biomarker for quantifying biological aging, typically estimated from structural MRI scans [11,5]. As illustrated in Fig. 1(a), brain age models are commonly trained on healthy individuals, and the difference between predicted brain age and chronological age, often referred to as the brain age delta, is used as an indicator of brain health. A positive delta is generally interpreted as accelerated aging [3,22]. However, recent studies suggest that brain age delta does not necessarily reflect accelerated aging, as brain age models built upon cross-sectional data primarily capture group-level averages of the training data and are not equipped to account for individual variability in aging rate [21,20]. For instance, an individual may consistently exhibit a positive delta over time, reflecting a stable baseline shift rather than an increased ongoing aging rate.

**Fig. 1.**
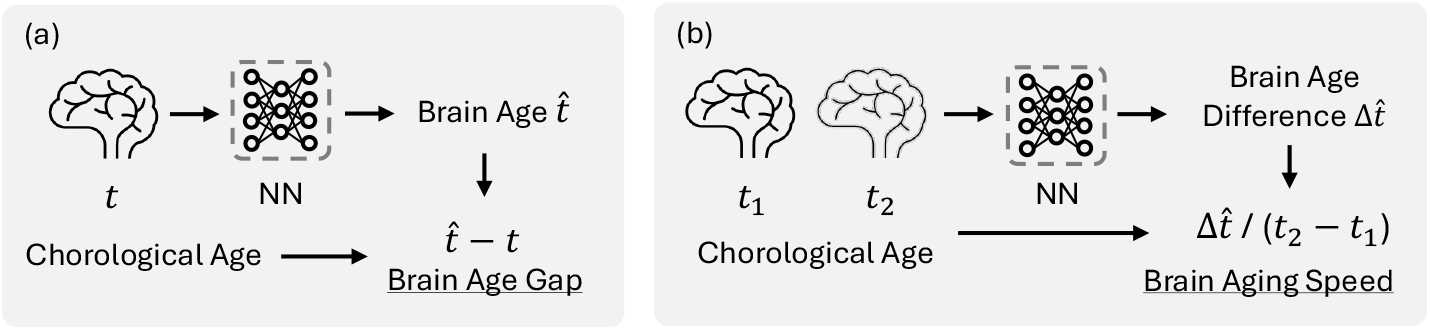
(a) Conventional cross-sectional brain age modeling predicts brain age from a single MRI scan. Brain age delta is defined as the difference between the predicted brain age and the chronological age. (b) Longitudinal brain aging modeling predicts the age difference between two MRI scans. Brain aging speed is computed as the predicted brain age difference normalized by the chronological time interval between scans.

Longitudinal studies provide a promising alternative for estimating aging dynamics by modeling temporal changes in brain imaging data. Recent approaches predict age differences directly from paired T1-weighted MRI scans [25,12], from which brain aging speed is computed as the ratio of the predicted age difference and the chronological time interval (Fig. 1(b)). However, existing methods primarily rely on intensity-based differences between scans, making them sensitive to imaging artifacts, scanner variability, and site effects. In addition, they typically yield a single global prediction, overlooking the spatial heterogeneity of brain aging patterns [14,7]. This can lead to suboptimal performance and reduce interpretability regarding region-specific contributions.

To overcome these limitations, we propose a novel framework that jointly characterizes local and global brain aging dynamics through morphology-driven modeling (Fig. 2). To derive representations more closely related to biological aging, we first estimate longitudinal morphological changes using diffeomorphic image registration, yielding a smooth, topology-preserving deformation field that describes anatomical transformations between scans. We then compute the Jacobian determinant of the deformation field to quantify voxel-wise expansion and contraction. To account for the spatial heterogeneity of aging dynamics, instead of producing a single global prediction, we partition the Jacobian map into non-overlapping patches and assign each patch to a dedicated local regressor to estimate regional brain age differences. These local predictions are subsequently integrated using learned patch relevance scores to generate a global age difference estimate. This design captures spatially varying aging patterns while enhancing both predictive performance and local interpretability.

**Fig. 2.**
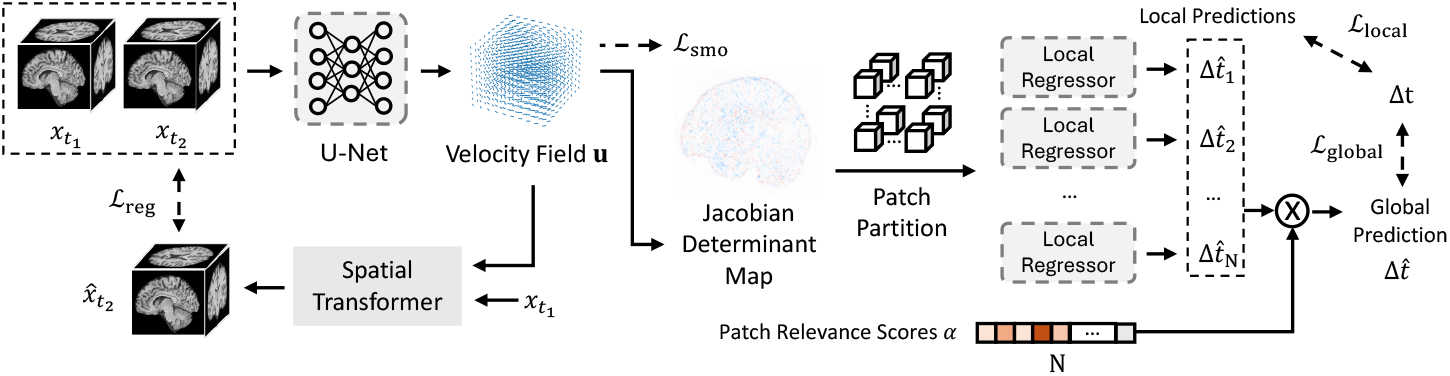
Overview of the proposed framework. Longitudinal morphological changes are first estimated using deep learning–based diffeomorphic image registration, producing a Jacobian determinant map that represents voxel-wise expansion or contraction. The Jacobian map is then partitioned into non-overlapping patches, and each patch is processed by a dedicated local regressor to predict regional aging speed. The local predictions are subsequently integrated using learned patch relevance scores, yielding a global prediction and an interpretable measure of each patch’s contribution.

Our contribution are as follows: 1) We propose a morphology-driven framework that estimates brain aging speed from registration-derived deformation fields; 2) We introduce a patch-wise regression strategy that jointly produces local and global predictions, accounting for spatial heterogeneity while enhancing interpretability; 3) We validate our method on a large and diverse dataset, demonstrating improved prediction performance and effectiveness in identifying abnormal aging patterns in diseased populations.

## 2 Method

In this section, we present the proposed aging speed estimation framework, as shown in Fig. 2. Given a pair of MRI scans, 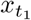 and 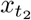, acquired from two visits of the same subject with chronological ages *t*_1_ *< t*_2_, our goal is to learn a mapping that predicts the age difference 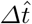 between the two scans. The framework is trained on cognitively normal subjects to capture normative aging dynamics. When applied to new data, the aging speed *v* is defined as the ratio between the predicted age difference and the actual chronological interval: 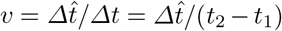. Intuitively, *v* = 1 corresponds to normal aging, and *v >* 1 indicates accelerated aging.

### Modeling morphological changes

We model longitudinal morphological changes between 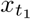 and 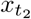 using diffeomorphic image registration that is implemented with a deep neural network [15]. Specifically, 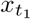 and 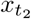 are concatenated and fed into a U-Net to predict a stationary velocity field *u* ∈ ℝ^3*×*H*×*W*×*D^, where H × W × D denotes the spatial dimensions of the input images. The diffeomorphic deformation field *ϕ*, which defines a smooth and topology-preserving transformation *ϕ* : ℝ^*H×W×D*^ → ℝ^*H×W×D*^ that warps 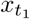 toward 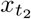, is obtained by integrating the ordinary differential equation parameterized by *u*:

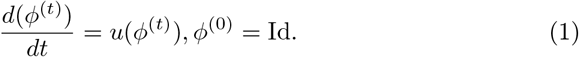

Integrating Eq. 1 over unit time yields the final deformation field:

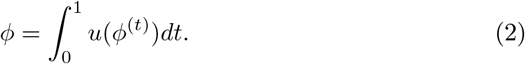

A spatial transformer layer [4] is then used to warp 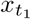 using *ϕ* to align it with 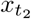. Registration accuracy is enforced using a local normalized cross-correlation loss [4] ℒ_*reg*_ between the warped image 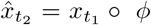 and 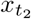. Following [6], a smoothness regularization term ℒ_*smo*_ is further applied to the velocity field *u* using a diffusion regularizer on its spatial gradients.

The deformation field describes voxel-wise displacement from 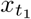 to 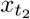, but does not directly quantify local volumetric changes. Therefore, we further compute the Jacobian determinant *J* ∈ ℝ^H*×*W*×*D^ of the deformation field *ϕ*, which characterizes voxel-wise expansion (*J >* 1) or contraction (*J <* 1). This Jacobian map serves as a morphology-based representation of longitudinal brain changes for subsequent aging speed estimation.

### Local predictions

Because aging is heterogeneous and different brain regions may exhibit distinct aging related patterns, we partition the Jacobian determinant map into N non-overlapping patches *J*_*i*_, *i* ∈ [1, N]. A patch-specific regression model is assigned to each patch to estimate the local brain age difference 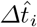. To encourage local predictions to approximate the true chronological age difference *Δt* = *t*_2_ − *t*_1_, we use the mean absolute error (MAE) loss:

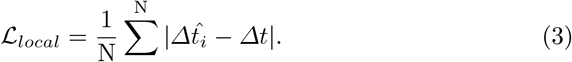

### Global prediction

The global prediction 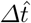 is obtained by aggregating local predictions using a set of patch relevance scores *α*_*i*_:

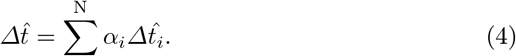

The relevance weights *α*_*i*_ are set as learnable parameters and are jointly optimized with other model parameters. To ensure non-negativity and unit summation, the Softmax function is applied to the relevance scores. The global prediction is also supervised with an mean absolute error (MAE) loss:

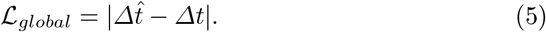

The final objective is the weighted sum of the registration loss, the smooth loss, the local prediction loss and the global prediction loss, where the weights are hyperparameters that control the strength of each term.

## 3 Experiments

**Datasets and participants**. Longitudinal T1-weighted MR images were collected from three cohorts: the Alzheimer’s Disease Neuroimaging Initiative (ADNI) [17], Open Access Series of Imaging Studies (OASIS-3) [13], and Australian Imaging Biomarkers and Lifestyle Study of Aging (AIBL) [8]. A total of 3915 imaging sessions from 1001 cognitively unimpaired (CU) subjects were used to train the proposed framework. Each subject had at least two sessions, and all sessions have valid age and diagnosis information. A subject was defined as CU if all of their sessions were consistently labeled as cognitively unimpaired. For every subject, all possible image pairs were constructed with chronological age differences between 1 and 16 years. The ADNI and OASIS-3 cohorts were randomly divided at the subject level into training (3087 sessions from 795 subjects, 5539 image pairs), validation (343 sessions from 94 subjects, 544 image pairs), and internal test sets (485 sessions from 112 subjects, 1091 image pairs). The AIBL cohort was used as an independent external test set (502 sessions from 155 subjects, 657 image pairs) to evaluate model generalizability. Subject-level splitting was performed to ensure that images from the same subject did not appear in multiple data partitions. The summary of the dataset can be found in Table 1.

**Table 1.**
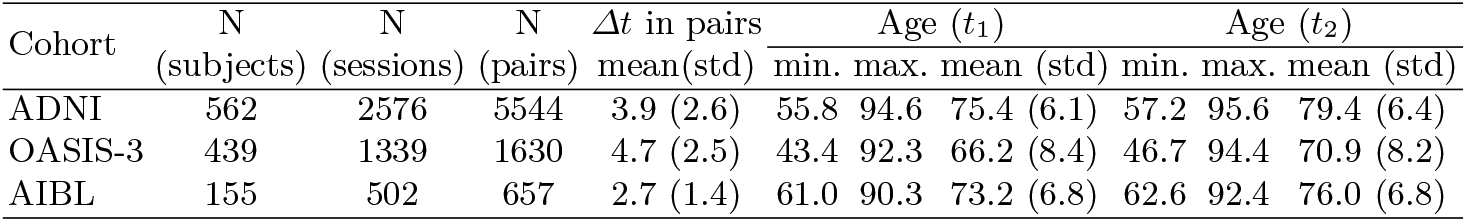
Summary of cognitive unimpaired participants used to train and validate the framework. The numbers of subjects, sessions, and image pairs are reported, together with the statistics of *Δt, t*_1_, and *t*_2_ (measured in years).

### MRI preprocessing

All MRIs were preprocessed using standardized steps, including bias field correction [24] and affine registration to align each image to MNI space [1,2] defined by the ICBM 2009c nonlinear symmetric template [10,9]. To reduce computational burden, the preprocssed images were cropped to remove background, resulting in a final image resolution of 176 × 192 × 176.

### Implementation details

We adopted the U-Net architecture from [18]. The local regressors were implemented using a lightweight CNN backbone [16]. Model training was performed using the Adam optimizer with a learning rate of 10^*−*3^ and a batch size of 8. The framework was trained for 100 epochs, requiring approximately 40 hours on an NVIDIA A40 GPU. To balance predictive performance and computational efficiency, the Jacobian determinant map was partitioned into 4 × 4 × 4 = 64 non-overlapping patches.

### Comparison with existing methods and ablation experiments

We compared the proposed method with: (1) SFCN [16], a widely used cross-sectional brain age prediction model, with the age differences computed as the difference between two individual predictions; (2) LM [25], which estimates age differences from voxel-level differences between paired scans; and (3) LILAC [12], which predicts age differences using feature-level differences between paired scans. In addition, two ablated versions of our framework were evaluated: (4) a variant that replaces the Jacobian determinant map with raw image differences while retaining the local prediction module (w/o morphological changes); and (5) a variant that performs global prediction only using the Jacobian determinant map without the local prediction module (w/o local prediction).

As shown in Table 2, longitudinal methods (LM, LILAC, and ours) substantially outperformed the cross-sectional approach (SFCN). Incorporating morphological information to predict temporal changes (Ours w/o local predictions) consistently improved performance on both internal and external test sets. Similarly, accounting for spatial heterogeneity via local predictions (Ours w/o morphological changes) also led to consistent gains. Integrating both components in the full framework further enhanced performance, highlighting the effectiveness of the proposed method.

**Table 2.** Performance comparison with existing methods and ablation models. Evaluation metrics include mean absolute error (MAE), root mean squared error (RMSE), and coefficient of determination (R^2^). Best performance is highlighted in **bold**.

| Methods | Longitudinal | Internal Test |  |  | External Test |  |  |
| --- | --- | --- | --- | --- | --- | --- | --- |
| | | MAE ↓ | RMSE ↓ | $R^2$ ↑ | MAE ↓ | RMSE ↓ | $R^2$ ↑ |
| SFCN [16] | ✗ | 2.35 | 3.21 | 0.26 | 1.41 | 1.79 | 0.26 |
| LM [25] | ✓ | 1.47 | 2.03 | 0.58 | 0.96 | 1.30 | 0.32 |
| LILAC [12] | ✓ | 1.66 | 2.77 | 0.36 | 0.96 | 1.29 | 0.51 |
| Ours (w/o morphological changes) | ✓ | 1.25 | 1.81 | 0.60 | 0.83 | 1.11 | 0.50 |
| Ours (w/o local predictions) | ✓ | 1.33 | 1.77 | 0.58 | 0.87 | 1.23 | 0.56 |
| Ours | ✓ | **1.13** | **1.59** | **0.68** | **0.70** | **0.94** | **0.60** |

### Effects of patch size

We evaluated the impact of patch size on model performance using three partition settings: 2 × 2 × 2 patches, 4 × 4 × 4 patches, and 6 × 6 × 6 patches, where the Jacobian determinant map is evenly divided along the three spatial dimensions. Smaller patch sizes generally provide finer-grained interpretability. However, experimental results (Table 3) show that excessively increasing the number of patches may degrade performance, likely due to the increased model complexity as each patch is associated with an independent local regressor. To balance interpretability, predictive performance, and computational efficiency, we select 4×4×4 patch partitions for subsequent experiments.

**Table 3.** Effects of patch size on prediction performance. Jacobian determinant map is evenly divided along the three spatial dimensions into specific number of patches. Best performance is highlighted in **bold**.

| Number of Patches | Internal Test |  |  | External Test |  |  |
| --- | --- | --- | --- | --- | --- | --- |
|  | MAE ↓ | RMSE ↓ | R<sup>2</sup> ↑ | MAE ↓ | RMSE ↓ | R<sup>2</sup> ↑ |
| 2 × 2 × 2 | **1.10** | **1.46** | 0.65 | 0.86 | 1.10 | 0.57 |
| 4 × 4 × 4 | 1.13 | 1.59 | **0.68** | **0.70** | **0.94** | **0.60** |
| 6 × 6 × 6 | 1.31 | 1.97 | 0.51 | 0.73 | 0.99 | 0.57 |

### Aging speed in disease populations

To validate the proposed method’s ability to model aging dynamics and the ability to detect abnormal aging, we applied the trained model to diagnostic transition groups defined by changes between the first and last visits, including two non-progression groups (CU-to-CU, MCI-to-MCI), and two progression groups (CU-to-MCI, MCI-to-AD).

Aging speed was computed as 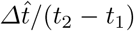, and the trend of average aging speed across different age groups is visualized in Fig. 3(a). For comparison, brain age delta was derived using SFCN predictions as a cross-sectional biomarker and is shown in Fig. 3(b). The non-progression groups (CU-to-CU and MCI-to-MCI) exhibit relatively constant aging speeds close to 1.0, suggesting stable aging dynamics. In contrast, the progression groups (CU-to-MCI and MCI-to-AD) show higher aging speeds, indicating accelerated aging. Meanwhile, while the CU-to-CU group has brain age delta values centered around 0, other groups involving diseased subjects tend to exhibit consistently positive brain age delta values. These observations suggest that brain age delta primarily reflects cumulative brain health status rather than ongoing aging rate, highlighting the advantage of modeling longitudinal aging dynamics.

**Fig. 3.**
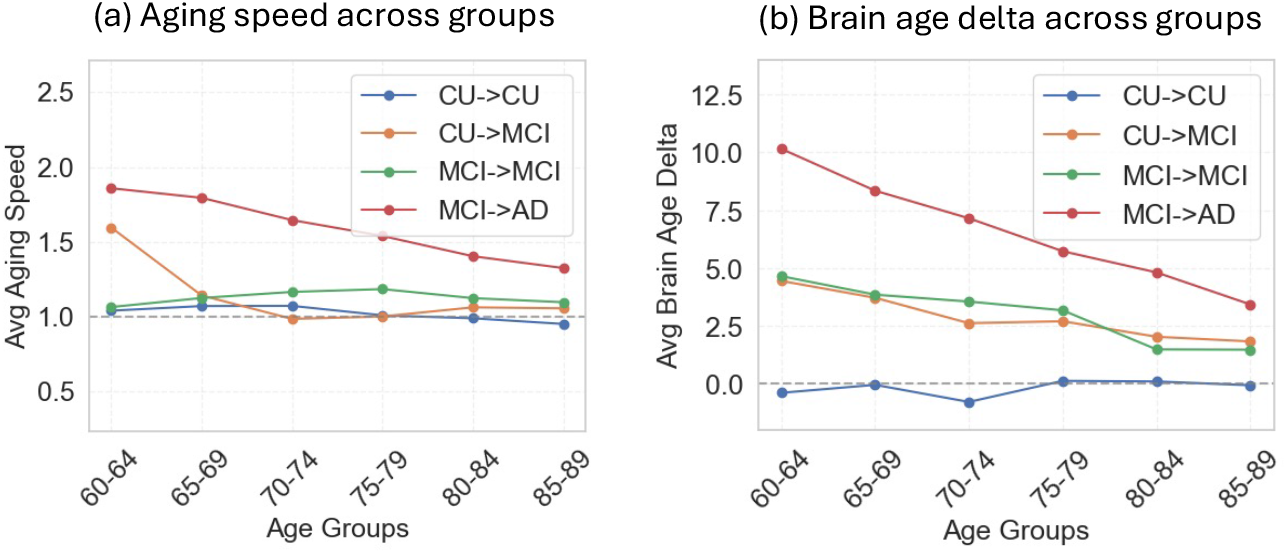
Trends of brain aging speed and brain age delta in different conversion groups. Subjects are binned into age groups, and the averages are calculated within each bin.

### Region contributions

We visualized the learned patch relevance scores and project them onto the ICBM 2009 brain atlas for better interpretability, as shown in Fig. 4. The model automatically learns sparse relevance patterns without explicit sparsity regularization, suggesting that only a subset of regions contributes substantially to aging speed prediction. In particular, subcortical regions appear to play an important role in modeling longitudinal aging dynamics.

**Fig. 4.**
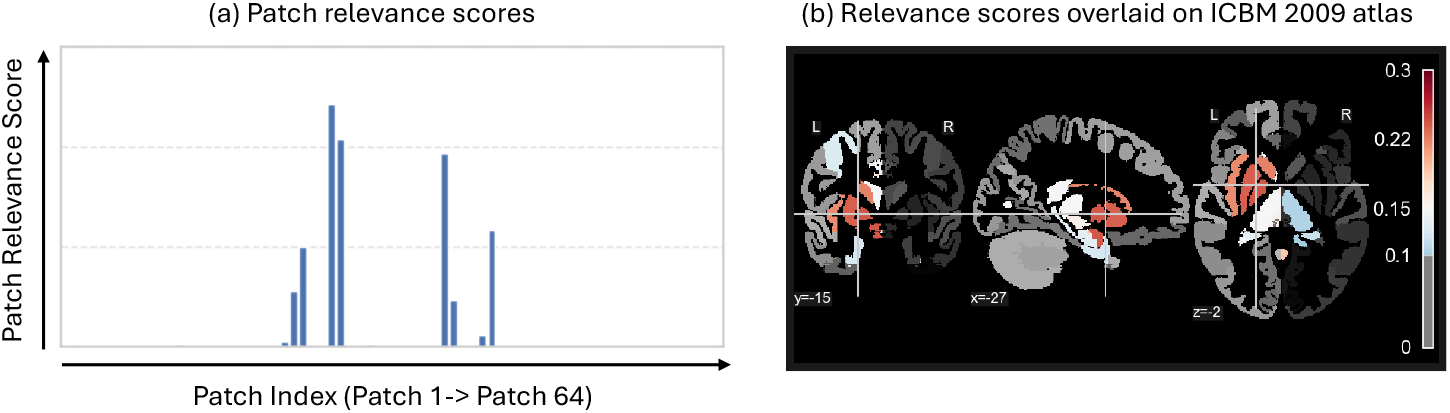
Visualization of learned patch relevance scores. (a) Distribution of patch relevance scores, where the x-axis represents patch indices and the y-axis represents relevance score values. (b) Brain region visualization obtained by mapping patch relevance scores onto the ICBM 2009 atlas. When a brain region overlaps with multiple patches, the regional score is computed as the average of the overlapping patch scores.

## 4 Conclusions

In this paper, we proposed a morphology-driven framework for estimating brain aging speed from longitudinal MRI scans. Unlike conventional brain age models that rely on cross-sectional and intensity-based representations, our method leverages diffeomorphic registration to capture anatomical deformation patterns and uses Jacobian determinant maps to characterize voxel-wise morphological changes. Additionally, by incorporating a patch-wise regression strategy, the framework jointly models local and global aging dynamics, enabling improved predictive performance while providing interpretability about regional contributions. Experimental results on large-scale longitudinal datasets demonstrate that the proposed approach outperforms existing cross-sectional and longitudinal methods, and can identify abnormal aging patterns in diseased populations.

## References

1. Avants, B.B., Epstein, C.L., Grossman, M., Gee, J.C.: Symmetric diffeomorphic image registration with cross-correlation: evaluating automated labeling of elderly and neurodegenerative brain. Medical image analysis 12(1), 26–41 (2008)

2. Avants, B.B., Tustison, N.J., Stauffer, M., Song, G., Wu, B., Gee, J.C.: The insight toolkit image registration framework. Frontiers in neuroinformatics 8, 44 (2014)

3. Baecker, L., Garcia-Dias, R., Vieira, S., Scarpazza, C., Mechelli, A.: Machine learning for brain age prediction: Introduction to methods and clinical applications. EBioMedicine 72 (2021)

4. Balakrishnan, G., Zhao, A., Sabuncu, M.R., Guttag, J., Dalca, A.V.: Voxelmorph: a learning framework for deformable medical image registration. IEEE transactions on medical imaging 38(8), 1788–1800 (2019)

5. Cole, J.H., Poudel, R.P., Tsagkrasoulis, D., Caan, M.W., Steves, C., Spector, T.D., Montana, G.: Predicting brain age with deep learning from raw imaging data results in a reliable and heritable biomarker. NeuroImage 163, 115–124 (2017)

6. Dalca, A.V., Balakrishnan, G., Guttag, J., Sabuncu, M.R.: Unsupervised learning of probabilistic diffeomorphic registration for images and surfaces. Medical image analysis 57, 226–236 (2019)

7. Eavani, H., Habes, M., Satterthwaite, T.D., An, Y., Hsieh, M.K., Honnorat, N., Erus, G., Doshi, J., Ferrucci, L., Beason-Held, L.L., et al.: Heterogeneity of structural and functional imaging patterns of advanced brain aging revealed via machine learning methods. Neurobiology of aging 71, 41–50 (2018)

8. Ellis, K.A., Bush, A.I., Darby, D., De Fazio, D., Foster, J., Hudson, P., et al.: The Australian imaging, biomarkers and lifestyle (AIBL) study of aging: methodology and baseline characteristics of 1112 individuals recruited for a longitudinal study of Alzheimer’s disease. International psychogeriatrics 21(4), 672–687 (2009)

9. Fonov, V., Evans, A.C., Botteron, K., Almli, C.R., McKinstry, R.C., Collins, D.L., Group, B.D.C., et al.: Unbiased average age-appropriate atlases for pediatric studies. Neuroimage 54(1), 313–327 (2011)

10. Fonov, V.S., Evans, A.C., McKinstry, R.C., Almli, C.R., Collins, D.: Unbiased nonlinear average age-appropriate brain templates from birth to adulthood. NeuroImage 47, S102 (2009)

11. Franke, K., Ziegler, G., Klöppel, S., Gaser, C., Initiative, A.D.N., et al.: Estimating the age of healthy subjects from T1-weighted MRI scans using kernel methods: exploring the influence of various parameters. Neuroimage 50(3), 883–892 (2010)

12. Kim, H., Karaman, B.K., Zhao, Q., Wang, A.Q., Sabuncu, M.R., Initiative, A.D.N.: Learning-based inference of longitudinal image changes: Applications in embryo development, wound healing, and aging brain. Proceedings of the National Academy of Sciences 122(8), e2411492122 (2025)

13. LaMontagne, P.J., Benzinger, T.L., Morris, J.C., Keefe, S., Hornbeck, R., Xiong, C., et al.: OASIS-3: longitudinal neuroimaging, clinical, and cognitive dataset for normal aging and alzheimer disease. MedRxiv pp. 2019–12 (2019)

14. Li, H., Cui, Z., Cieslak, M., Salo, T., Moore, T.M., Gur, R.E., Gur, R.C., Shinohara, R.T., Oathes, D.J., Davatzikos, C., et al.: Spatial heterogeneity and subtypes of functional connectivity development in youth. Nature Communications (2026)

15. Li, H., Fan, Y., Initiative, A.D.N.: Mdreg-net: Multi-resolution diffeomorphic image registration using fully convolutional networks with deep self-supervision. Human Brain Mapping 43(7), 2218–2231 (2022)

16. Peng, H., Gong, W., Beckmann, C.F., Vedaldi, A., Smith, S.M.: Accurate brain age prediction with lightweight deep neural networks. Medical image analysis 68, 101871 (2021)

17. Petersen, R.C., Aisen, P.S., Beckett, L.A., Donohue, M.C., Gamst, A.C., Harvey, D.J., Jack Jr, C.R., Jagust, W.J., Shaw, L.M., Toga, A.W., et al.: Alzheimer’s disease neuroimaging initiative (ADNI) clinical characterization. Neurology 74(3), 201–209 (2010)

18. Pombo, G., Gray, R., Cardoso, M.J., Ourselin, S., Rees, G., Ashburner, J., Nachev, P.: Equitable modelling of brain imaging by counterfactual augmentation with morphologically constrained 3D deep generative models. Medical Image Analysis 84, 102723 (2023)

19. Raz, N., Rodrigue, K.M.: Differential aging of the brain: patterns, cognitive correlates and modifiers. Neuroscience & Biobehavioral Reviews 30(6), 730–748 (2006)

20. Smith, S.M., Miller, K.L., Nichols, T.E.: Characterising ongoing brain aging and baseline effects from cross-sectional data. Imaging Neuroscience 3, IMAG–a (2025)

21. Smith, S.M., Vidaurre, D., Alfaro-Almagro, F., Nichols, T.E., Miller, K.L.: Estimation of brain age delta from brain imaging. Neuroimage 200, 528–539 (2019)

22. Tanveer, M., Ganaie, M., Beheshti, I., Goel, T., Ahmad, N., Lai, K.T., Huang, K., Zhang, Y.D., Del Ser, J., Lin, C.T.: Deep learning for brain age estimation: A systematic review. Information Fusion 96, 130–143 (2023)

23. Toepper, M.: Dissociating normal aging from Alzheimer’s disease: A view from cognitive neuroscience. Journal of Alzheimer’s disease 57(2), 331–352 (2017)

24. Tustison, N.J., Avants, B.B., Cook, P.A., Zheng, Y., Egan, A., Yushkevich, P.A., Gee, J.C.: N4ITK: improved N3 bias correction. IEEE transactions on medical imaging 29(6), 1310–1320 (2010)

25. Yin, C., Imms, P., Chowdhury, N.F., Chaudhari, N.N., Ping, H., Wang, H., Bogdan, P., Irimia, A., Initiative, A.D.N.: Deep learning to quantify the pace of brain aging in relation to neurocognitive changes. Proceedings of the National Academy of Sciences 122(10), e2413442122 (2025)

